# The Kinetic Intron Hypothesis

**DOI:** 10.64898/2026.03.04.709683

**Authors:** Garrett Tisdale

## Abstract

Intron length is a fascinating example of form without function. The vast majority of intronic space within genomes remains without a provided utility. It often fascinates us to find introns performing any function at all, establishing an attention bias against the vast lacking of utility of the remaining intergenic space. In an attempt to better understand the greater breadth of intronic length, I investigate here what I term The Kinetic Intron Hypothesis. This hypothesis investigates hypothetical dynamics of intron RNA synthesis and degradation. It explores how NTPs stored within intron RNA might function in mitosis and NTP resource management. Preliminary testing of the hypothesis leads to trends that warrant further exploration and validation by the scientific community.

**Significance:** Currently no widely acknowledged model exists to characterize the length of introns within genes, yet intron length is massively abundant in eukaryotic genomes. Here I present an attempt to model the length of introns. In doing so, I explore novel hypothesized intron dynamics, presenting preliminary data for previously uncharacterized intron characteristics. The new data and model have the protentional to unveil new avenues of utility for introns at the intracellular level.

## Introduction

The length of introns within eukaryotes remains a perplexing enigma. As discrete units—whether through alternative splicing, nonsense mediated decay sites, or intron delay—introns appear to serve within core biological pathways. Inversely, the continuously variable length of introns within genomes is seemingly random and uncorrelated with a perceived function. Within the human genome, 87.9% of transcribed genetic material is intronic (1). I take this as an unexplained absurdity. My fascination can be simply stated: What constitutes the length of introns within eukaryotes?

The dominance of introns within genomes originates from the last eukaryotic common ancestor (LECA) (2). Phylogenetic analysis signifies that this widespread incorporation of introns is a product of eukaryogenesis. The general view for the mechanism of intron incorporation into LECA’s genome consists of a α-proteobacteria containing group II introns being engulfed and incorporated though endosymbiosis. Intron proliferation after that point was dramatic and unprecedented. Koonin elaborates that intron proliferation could have reached ≥2 introns per kbp and >70% of the protoeukaryotic genome (3). Following the explosion of introns in pre-LECA, a reduction in introns occurred. Much study has suggested that by the time of LECA spliceosome systems and machinery had already become established thereby solidifying the core mechanisms for spliceosomal introns within all eukaryotes.

Despite the reasonably projected origins and known mechanisms by which transcribed introns are synthesized and degraded, no unifying principle underlies their existence. Even within modern reviews authors still conclude, “The existence of introns in [the] genome is a real mystery” (4). Specific functions have given meaning to select introns, but these functions do not describe an essential essence for which the prevalence of all intron length is deemed necessary within eukaryotes. The Synthetic Yeast Genome Project, in efforts to streamline *Saccharomyces cerevisiae*, removed many introns with no overt phenotypic consequences under standard laboratory conditions (5). These findings have contributed to ongoing debate regarding whether introns provide broadly essential functions or instead confer primarily context-dependent regulatory advantages. However, the observation that introns comprise approximately 87.9% of transcribed genetic material is difficult to reconcile with a model in which their utility is predominantly context-dependent. This strongly suggests the existence of more general and fundamental roles underlying their pervasive abundance.

Without a basis by which to begin explaining intron length, I speculated that some unknown observable has yet to be investigated. One hypothesis relating to intron RNA kinetics piqued my interest. I was curious if the length of introns and their synthesis/degradation kinetics could modulate intracellular NTP levels, noting that introns are incredibly long, continuously transcribed, and take up a considerable resource burden for cells. As a steady state model this hypothesis quickly failed. The rapid turnover of intron lariats leaves the pool of introns indistinguishable from the starting pool of NTPs. Yet the model did produce an equation reminiscent of data produced in 2002 by Castillo Davis et al. (6), where it was shown that intron length is roughly inversely proportional to gene expression (replicated for mouse in Figure S4A). While the initial model was wrong, something about the model resembled the literature data and was therefore of interest for further exploration.

As a kinetics problem, the paradigm of rapid intronic RNA degradation left the steady state kinetic-intron model untenable as a justification for intron length. As such I searched for potential scenarios where intron lariat degradation may be halted. Notably it is established as a paradigm that the degradation of intronic mRNAs is exceedingly rapid. Experimentally, intron probes for single molecular fluorescence in situ hybridization (smFISH) are used for identifying transcription sites, as individually spliced introns are degraded so rapidly after transcription that they do not appear elsewhere but their place of origin. If the paradigm of rapid intron degradation were to be broken, I posited that the inverse relationship between intron length and gene expression could hypothetically be explained via intron kinetics. Only two plausible scenarios came to mind: 1.) Preserving intronic RNA for a rapidly accessible NTP reservoir for kick starting transcription post starvation, when the pentose phosphate pathway is shut down or 2.) Preserving intronic RNA for a rapidly accessible pool of NTPs post mitosis for the rapid synthesis of the G1 transcriptome after mitotic transcriptional silencing. Both are scenarios where a discontinuous break in the kinetics of the central dogma may require an imbalance of NTP resources. Mitosis kinetics was easier to explore. This sporadic rationale led me to investigate the relationship between three key associations: intron length, gene expression kinetics, and mitosis. The hypothetical model underlying the relationship between the hypothetical kinetics of intron preservation and degradation during mitosis is presented in the supplemental and is termed ‘The Kinetic Intron Hypothesis’.

## Preliminary Results

### Total Intronic Length Density is Greatest in Genes Overexpressed During Mitosis

The Kinetic Intron Hypothesis predicts introns as a core functional component of mitosis; therefore, I began by analyzing the distribution of total intronic length in genes overexpressed during mitosis. Tanenbaum et al. captured RNASeq data of the G2, M, and G1 transcriptomic states in non-transformed human epithelial cells (7). Expression values in the form of reads per kilobase per million hits (RPKM) values from this data were matched with the total summed intronic length within corresponded genes using the Hg19 genome (8) and compared across several folds of upregulation (Table S1). Early during the analysis, it was noted that a group of genes with extremely large outlier introns (those with total intronic length greater than 100kbp) greatly impacted the mean total intronic length while establishing large standard deviations. On average this consisted of only ~10% of the data (Fig 1 E and F). As such the data was bifurcated such that most introns, those below 100k bp, could be examined separately from the extremes.

**Figure 1:**
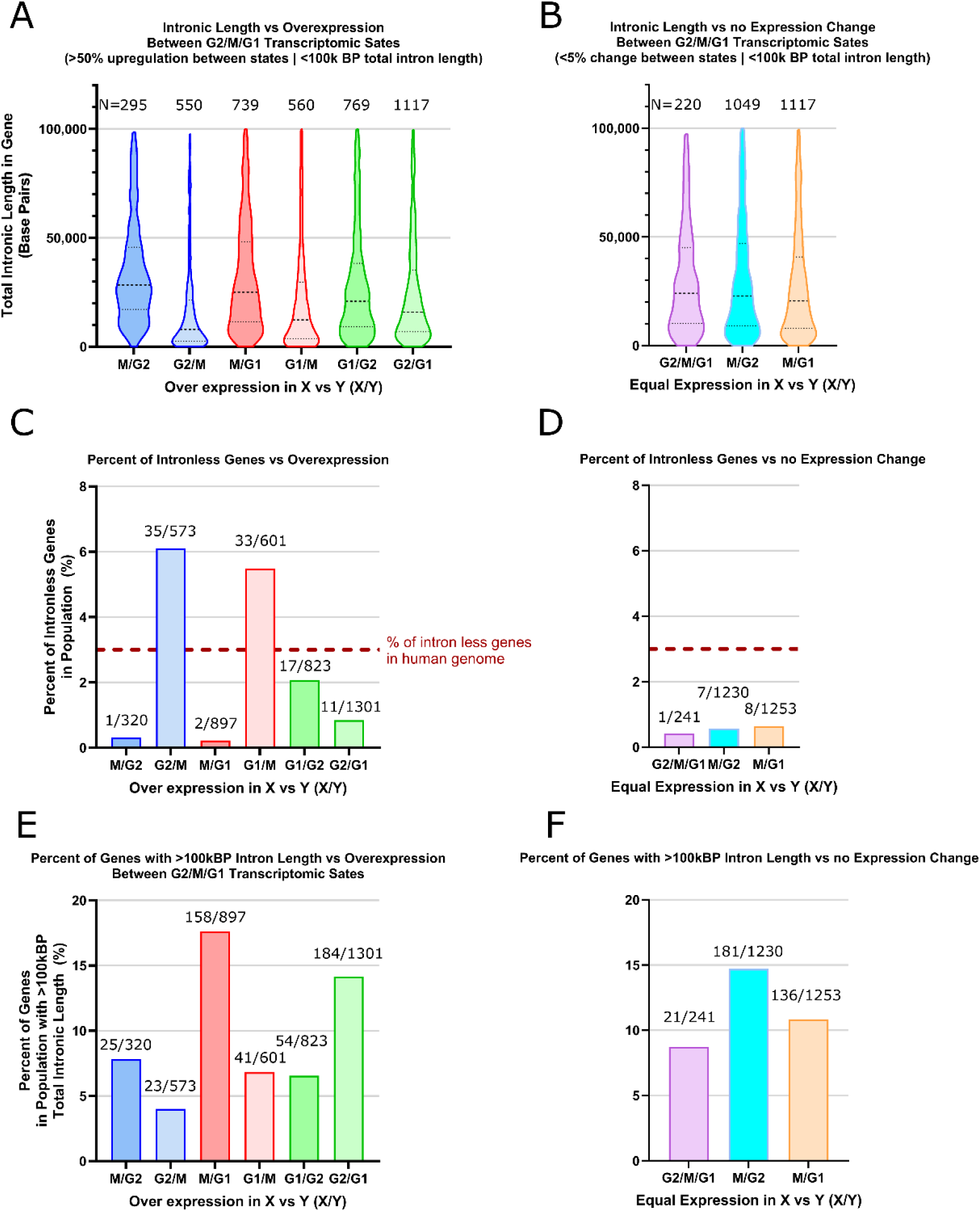
(A,B) Violin plots showing the distribution of total intron length among genes shorter than 100 kbp that are upregulated by more than 50% in state X relative to state Y (X/Y). (C,D) Percentage of intron-less genes among genes exhibiting greater than 50% upregulation between cell-cycle states. (E,F) Percentage of genes longer than 100 kbp among genes exhibiting greater than 50% upregulation between cell-cycle states. **Gene expression data were obtained from Tanenbaum et al. (7), and intron annotations were derived from the hg19 human genome annotation (8). All analyses and integration of these datasets were performed in this study.

Examining distributions in Table S1 and Figure 1 reveals a plethora of information. 1.) Genes with an increase of more than 50% in expression during mitosis contain approximately twice the total intronic density as compared to genes overexpressed in G1 and G2. 2.) When examining genes with >50% increase in expression during mitosis there is only 1 reported intron-less gene in the M/G2 group and 2 in the M/G1 group (<1% of genes), indicating almost all genes overexpressed during mitosis contain introns (Figure 2C). Opposite therein, genes under expressed during mitosis as compared to G2 and G1 are vastly overrepresented by intron-less genes (6.11% and 5.49% respectively) compared to the whole genome statistic of ~3% (1). 3.) Most genes do not greatly change their transcriptomic state between G2, M, and G1 especially when comparing the transition between G2 and M.

**Figure 2:**
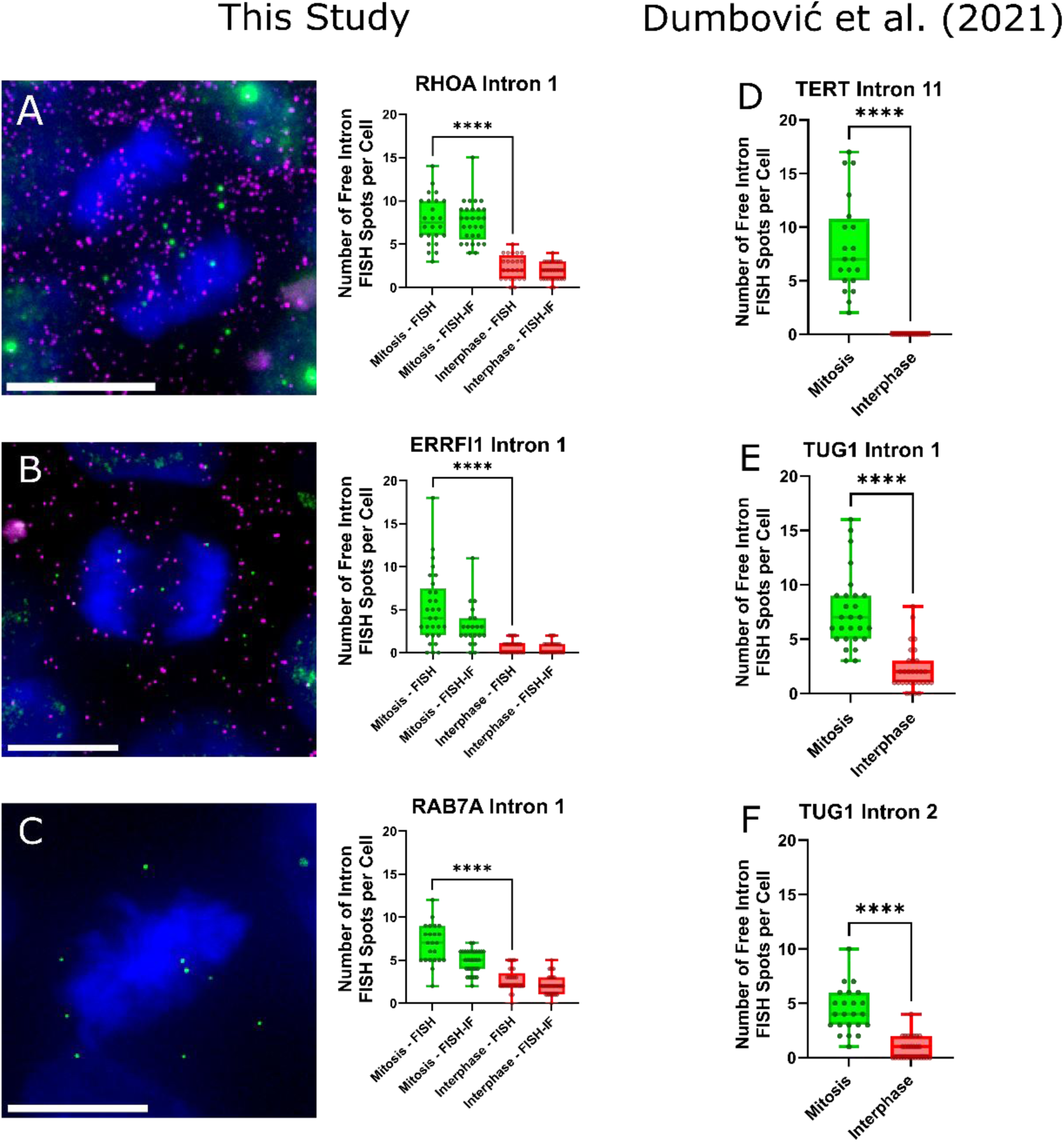
Free (spliced) intron RNA FISH spots (green) persist into mitosis and are delocalized from exon FISH spots (magenta) in human induced pluripotent stem cells. Representative examples are shown for *RHOA* intron 1 (A) and *ERRFI1* intron 1 (B). Intron RNA FISH spots accumulate during mitosis (C; exon probe not acquired to validate splicing state). **Panels D–F show data originally reported by Dumbović et al. (12), which were reanalyzed and re-plotted in this study under CC by 4.0 (https://creativecommons.org/licenses/by/4.0/). Scale bar = 10 µm. **** p ≤ 0.0005.

Examination of the violin plots of total gene-wise intron length distribution across the different upregulated states reveals a clear distribution change between M/G2 and G2/M (Figure 2A). While the G2/M upregulated state clusters near zero, its M/G2 counterpart appears to have an even distribution of intron density between ~10kBP and ~40kBP. This distribution persists, but is less prominent, for the M/G1 upregulated state. As an internal control, comparisons between G2 and G1 are provided where the unique mitotic distributions appear less defined. Notably, the difference between the M and G2 transcriptomic states are small; genes expressed in G2 are also expressed leading into M. Trends occurring in M likely also contribute towards the trends occurring in G2 when compared against G1. When examining genes which do not change between the states G2, M, and G1, G2 and M, or G1 and M (<5% change), the distributions follow the over expressed mitotic states rather than the reduced intron density distributions in genes over expressed in G1 and G2 states when compared against M. The percentage of intron-less genes in those states likewise follows the same trend (Figure 2C). A clear bias against intronic density in genes under expressed during mitosis is observed.

### Intron RNA Persists into Mitosis

The most controversial aspect of The Kinetic Intron Hypothesis under examination is the breaking of the central dogma paradigm that intron RNAs are always rapidly degraded. Aside from a hand full of circular RNA of various functions (9) and a subgroup of introns in S. cerevisiae (10), no wide scale observation of intron RNA retention has been observed to my knowledge. Therefore, it is of the utmost importance to the validity of this hypothesis to test if spliced introns are preserved—or rather persist—during mitosis, opposite of what the paradigm would suggest. To test this hypothesis RNA smFISH was employed.

Three intron probes, and two corresponding exon probes were obtained. The intron/exon probes for the genes *ERRFI1* and *RHOA* and a intron only probe set for *RAB7A* had been prior validated for control samples in another publication (11). The three genes vary in their magnitude of expression, however, the G2/M/G1 data examined prior indicates that there is no noteworthy change in expression between the mitotic, G1, and G2 states for any of the genes (7). I acquired human induced pluripotent stem cells (iPS cells | CS25i-18n2), performed FISH, and mounted with a DAPI infused mounting media to visualize dividing nuclei. FISH-immunofluorescence (FISH-IF) staining was performed against E-cadherin to delineate cell boundaries within iPS cell colonies and examine the distribution of free introns in daughter cells (Figure S3). During FISH-IF, the original e-coil tRNA hybridization blocking reagent used in the FISH-only experiments was no longer available and was replaced with salmon sperm DNA. This substitution resulted in elevated background fluorescence and reduced RNA detection, consistent with decreased effective hybridization sensitivity. The attenuation was most pronounced for the *EEF1A1* and *RAB7A* intron probe sets (Figure 2 and Figure S3). Despite the diminished signal-to-noise ratio, the qualitative biological phenomena were reproducible.

All three genes revealed an increase in free intron accumulation, or simply intron accumulation for *RAB7A*, during mitosis as compared to their interphase controls (Figure 2). Note that in the interphase condition, suspected transcription sites were counted as intron spots due to a lack of discernability, thereby resulting in a likely over detection. This lack of discernability arose from the distance between the 5’ intron probes and 3’ exon probes; finished 3’ exons did not colocalize as they finished transcribing and quickly diffuse away. This experiment acted to expand upon recently published data by Dumbović et al. (12), as such their data is further shown for reference (Figure 2D-3F).

Two of the introns were of particular interest. Intron 11 of *TERT*, as studied by Dumbović et al., happens to only be spliced during mitosis, and interestingly the number of freely spliced introns correlates with the number of spliced mRNA (12). The other interesting gene, *ERRFI1*, possessed a very broad distribution of freely spliced introns during mitosis ranging from 0 to 18. In the study published by Wan et al. (11) *ERRFI1* showed drastically different transcriptional bursting dynamics than *RHOA* and *RAB7A*, possessing drastically longer “off” times but similar “on” times. The large variance in RNA can be reasonably deduced as arising from catching the bursting state in either the “on” or “off” state in the leads up to mitosis. Furthermore, the E-Cadherin IF revealed what appeared to be cells in late-stage cytokinesis at the end of their mitotic stage, each with lesser compacted DNA and with defined individual cell membranes. The cells further lacked apparent transcription sites and their DAPI stain was more dense and compact than interphase cells. Since it could not be reliably concluded that each ‘daughter’ cell was not a cell entering mitosis as opposed to exiting mitosis these cells were dropped from the analysis in Figure 2, however, the free intron spots were seen approximately split evenly between daughter cells. This suggests the persistence of spliced intron may exist as far into mitosis as late stage cytokinesis and distribute introns randomly among daughter cells.

With these three new introns and genes added to the list, the total number of reported genes to contain introns persisting into mitosis is seven. Prior to these data, a study examining labeled introns in drosophila embryos reported a buildup of introns during mitosis in the Delta gene (13). While it could be that authors simply do not report an absence of observation of intron RNA accumulation in mitotic cells, the seven reported so far contain a 100% success rate of persisting into mitosis. It should also be noted that free introns can be seen in the background of published FISH images (14, 15). Visualizing mitotic cells is also generally technically challenging. During nuclear envelope breakdown cells often bulge and become loosely attached to coverslips—often termed mitotic shake-off—thereby causing data loss of the mitotic condition. Such is not the case with iPS cells which grow in colonies.

## Discussion

Introns remain fascinating, and the characteristics explored here add to their mysteries. The data presented is not encompassing or exhaustive due to fiscal and other limitations, yet it reveals a potential novel link between intron length, gene expression, intron turnover kinetics, and mitosis.

Strange ideas can lead to strange data. The relative success of the model presented in the supplemental, and the oddly circumspect discoveries the model directed attention towards in the above figures, leads to a new set of creative hypotheses to continue the scientific method. Taken together, the data defines both motive and means: evolutionary pressure favors intron length as a dominant feature of mitotically expressed genes (Figure 1), while the persistence of intron lariats into mitosis provides a plausible mechanistic substrate (Figure 2). The remaining open question is opportunity—what functional process, if any, harnesses these properties during mitosis.

To the curious reader, I would direct attention to the suspicious dynamics of intron lariat turnover during mitosis. As noted, a general lack of intron turnover has yet to be seen, thereby making the preliminary sightings in Figure 2 one of key interest. If the RNA material of introns lacks a physical presence due to their rapid degradation and its absence therein, intron RNA further lacks the ability to be of any prolonged or abundant biophysical utility. The mere observation that intron turnover may be halted during mitosis opens a potential interpretation of reality where intron RNA may possess unique dynamic regulations and interactions. The added correlation with intron length and mitotic gene expression (Figure 1) significantly amplifies the peculiarity of the observation. Such potential biophysical mechanisms may fall within or beyond the scope of the model that predicted the behavior. Many other aspects of interest can be explored at depth in the supplemental.

To the degree that The Kinetic Intron Hypothesis and model is accurate, judgment is deliberately deferred. The emphasis of this work is not on the model itself, but on the novel characteristics it helped bring into focus. Indeed, the hypothetical model may be incorrect, incomplete, or entirely misguided. Yet ideas need not be fully accepted to be meaningfully explored. In this spirit, I invoke Schrödinger’s reflection: “I can see no other escape from this dilemma (lest our true aim be lost for ever) than that some of us should venture to embark on a synthesis of facts and theories, albeit with second-hand and incomplete knowledge of some of them and at the risk of making fools of ourselves” (16).

## Methods

### For Mus musculus (mouse) and Homo sapien (human)

BSgenome.Mmusculus.UCSC.mm10 (17) and TxDb.Hsapiens.UCSC.hg19.knownGene (8) R packages were used respectively in conjunction with the GenomicFeatures R package (18). Due to the presence of many alternatively spliced transcripts within the genomes of these two species the mean intron length and mean transcript/mRNA transcript lengths (containing the CDS lengths, 5’ UTR length, and 3’ UTR regions) were used for each gene as specific information of the alternative splicing of the transcripts was not provided. Mouse fibroblast kinetics data from Schwanhäusser (19) was matched to intron and exon values. For plotting and calculating Equation 1, the data were parsed of all NA values, 0 length intron values, and 0 transcription rates. Furthermore, one data point was excluded as an outlier from the transcript length profile of the mouse data as the outlier was orders of magnitude removed from the population.

For mouse calculations using Equation 1, each gene-wise calculation utilized the gene’s total gene-wise intronic length, transcript length, and mRNA steady state value with the duration of between the G2/M checkpoint (τ) estimated as 55 minutes (as described in the supplemental text). Binning was performed along the prediction axis from larger prediction values to lower prediction values, preserving the variance towards the extreme and dropping the remainder bin of values nearest 0 which was always more well behaved. The remainder was dropped to preserve equal variance between bins.

Human G2, M, and G1 expression data from Tanenbaum et al. (7) was matched to their corresponding transcript and total intron length as previously described. Fold change was determined as the ratio between any two states from RPKM measurements (RPKM_M_/ RPKM_G1_, RPKM_M_/ RPKM_G2,_ …). As performed by Tanenbaum et al., genes containing less than 200 total reads and more than threefold change between replicate conditions was further dropped during analysis. All graphs in Figures 1 and 2 were generated in Prism.

The standard FISH protocol was performed. 18mm #1 coverslips (Electron Microscopy Services, 72292-09) were washed in 3M sodium hydroxide (MilliporeSigma,221465) for 3 minutes prior to Geltrex coating (Thermos Fisher, A121330) diluted 1:100 in DMEM/F12 (Thermo Fisher, 12634010). Cs25i iPS cell were cultured in StemFlex media until fixation. Cells were washed twice with 1x phosphate buffered saline (PBS, Corning, 46-013CM) diluted with nuclease free water (Quality Biological, 351-029-131CS) and 5mM magnesium chloride (MilliporeSigma, M2670-500G) (together being PBSM) before fixing with 4% paraformaldehyde for 10 minutes. Cells were washed twice with PBSM with the last wash lasting for 10 minutes. Permeabilization followed for 10 minutes with 5% Triton-X100 (MilliporeSigma, T8787-100mL) diluted in PBMS. Cells were washed twice with PBSM with the last wash lasting for 10 minutes. Pre-hybridization buffer consisting of 2xSSC (saline-sodium citrate buffer, Corning, 46-020-CM), 10% formamide (Millipore Sigma, F9037-100ML), diluted in nuclease free water was allowed to incubate for 10 minutes. Cells were hybridized at 37°C for 3 hours in 10% formamide, 1mg/mL competitor E. coli tRNA (Millipore Sigma, 10109541001), 10% w/v dextran sulfate (Millipore Sigma, D8906-100G), 0.2 mg /mL BSA (VWR,0332-25G), 2x SSC, 10 units/mL SUPERaseIn (Thermo Fisher, AM2694), 100nM intron and exon probes, diluted in nuclease free water. Following hybridization cells were washed with 2x SSC for 2 hours and 21 minutes followed by two 10 minutes washes at 37°C with 1x phosphate buffered saline (PBS, Corning, 46-013CM). Cells were mounted on slides (Thermo Fisher, 12-552-3) with ProLong Diamond antifade reagent with DAPI (Invitrogen, P36962).

FISH-IF followed the FISH protocol with modifications. Permeabilization and pre-hybridization contained 5mg/mL of BSA for blocking (VWR, 0332-25G) and Superase RNAse inhibitor. Hybridization buffer contained at 0.2mg/mL UltraPure BSA (Invitrogen, AM2618), mouse E-Cadherin at 1:1000 dilution (BD Biosciences, 610182), and 0.1mg/mL salmon sperm in place of depleted stock of discontinued E. coli tRNA. Cells were washed four times for 5 minute durations using 2x SSC and 10% formamide at 37 °C. Then secondary antibody Cy7 labeled goat anti-mouse (Invitrogen, A-21037) at 1:1000 dilution was incubated twice at 37 °C for 20 minutes in 2x SSC and 10% formamide. Cells were washed with 2x SSC at room temperature four times at 5 minute durations prior to mounting in ProLong Diamond antifade reagent with DAPI.

Imagining was performed on a custom up-right Nikon Eclipse Ni-E microscope equipped with 60x oil immersion objective lens (1.4 NA, Nikon), Spectra X LED light engine (Lumencor), and Orca 4.0 v2 sCMOScamera (Hamamatsu). The x-y pixel size was 108.3 nm with 300 nm z-pixel size. Images were acquired with Nikon Elements.

Quantification of FISH images was performed manually in ImageJ/FIJI to account for the 3D and overlapping nature/structure of iPS cell colony formation as well as to effectively discriminate for the poor background labeling of the FISH probes in some samples. Colocalization was defined as two distinguishable diffraction limited spots within 300nm peak-to-peak. Diffraction limited 0.1um TetraSpek Microsphere (ThermoFisher Scientific, T7279) were seeded onto coverslips and imaged to evaluate chromatic aberration occurring between imaging conditions during colocalization. During one imaging session there was infrequent skipping of frames resulting in minor data loss. It was determined that the loss of data did not significantly influence the discernability of the data.

## Acknowledgements

I thank Dr. Daniel Larson for providing the FISH probes used to assess spliced intron RNA persistence during mitosis, as well as for his constructive discussions. I thank Dr. Pan Li for providing the iPSC lines used in these experiments, and Dr. Bin Wu for providing the facilities and microscopy resources required to complete the FISH experiments. I also thank Dr. Matthew Ferguson for his early support and valuable discussions throughout the course of this work. Finally, I thank Dr. Gamid Abatchev and Dr. Daniel Fologea for their assistance in revising the manuscript.

## Data and Code Availability

Analysis code and example FISH/FISH-IF images used during analysis are available at: https://drive.google.com/drive/folders/1rT_KonGHGtawlSYYJnEgUVWjt-yqOUvZ?usp=sharing. Complete FISH and FISH-IF images available upon request.

## Funding

This work was supported by the National Institutes of Health (NIH) under Institutional Training Grant [T32-GM135131]. The content is solely the responsibility of the author and does not necessarily represent the official views of the National Institutes of Health.

## Supplemental

### Model

The following model describes the dynamics of introns under the hypothesis that intron lariat reserve promotes a reservoir of NTPs. Specifically, this reservoir of NTPs is predicted to occur during the lead up to mitosis as the G2/M checkpoint promotes the transcriptomic state of mitosis prior to prophase. Following mitosis, the model promotes the idea that the reserved pool of NTPs establishes a reservoir of resources for the genesis of the G1 transcriptomic state. The necessity of the pool would be predicated on the assumption that simply elevating the existing pool of NTPs would result in homeostatic dysregulation of core metabolic processes via le Chatelier’s principle. It is also predicated on the assumption that the degradation of the G2 transcriptome does not directly feed NTPs to the G1 transcriptome (discussed in the literature review later). The intron reservoir would therefore act as a counterbalance to the NTP flux occurring during early G1 transcriptome synthesis (Supplemental Figure 1).

First it is established that a reserved pool of ribonucleotides would consist of a total number of NTPs (*N*). During the processes of intron lariat preservation, *N* NTPs would thus be divisible across the sum length of all introns (*i*) of a given population of genes (*g*) synthesized at a rate (*r*) over the duration of intron lariat reserve (τ) such that, *N* = Σ *i_g_* · *r_g_* · τ. Clearly note that from here on the division of the total length of all introns over individual introns within a gene is arbitrary and only the total sum length of all introns within a gene together represents *i*. Continuing, since *N* represents the total number of mono-ribonucleotides required to promote the G1 transcriptomic state, *N* may be equated to the sum of all mono-ribonucleotides composing that state such that 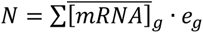 where 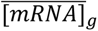 is the G1 mRNA steady state concentration of a given gene *g* and *e* is the mRNA transcript length of said gene. The equality is thus established, 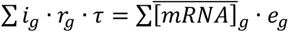. It should be noted that in ideal steady state 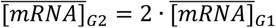 due to *r_G_*_2_ containing twice the number of transcribing loci per gene and thus twice the rate. However, each locus would be producing its equivalent product for its resulting daughter cell after division and is thus normalized. Furthermore, this equation explores the hypothetical global effect of the phenomena as though in temporal order. Gene-wise, however, the pressure exerted onto individual genes must exist irrelevant of the temporal dynamics between the mitotic and the G1 despite being composed of those variables.

**Supplemental Figure 1:**
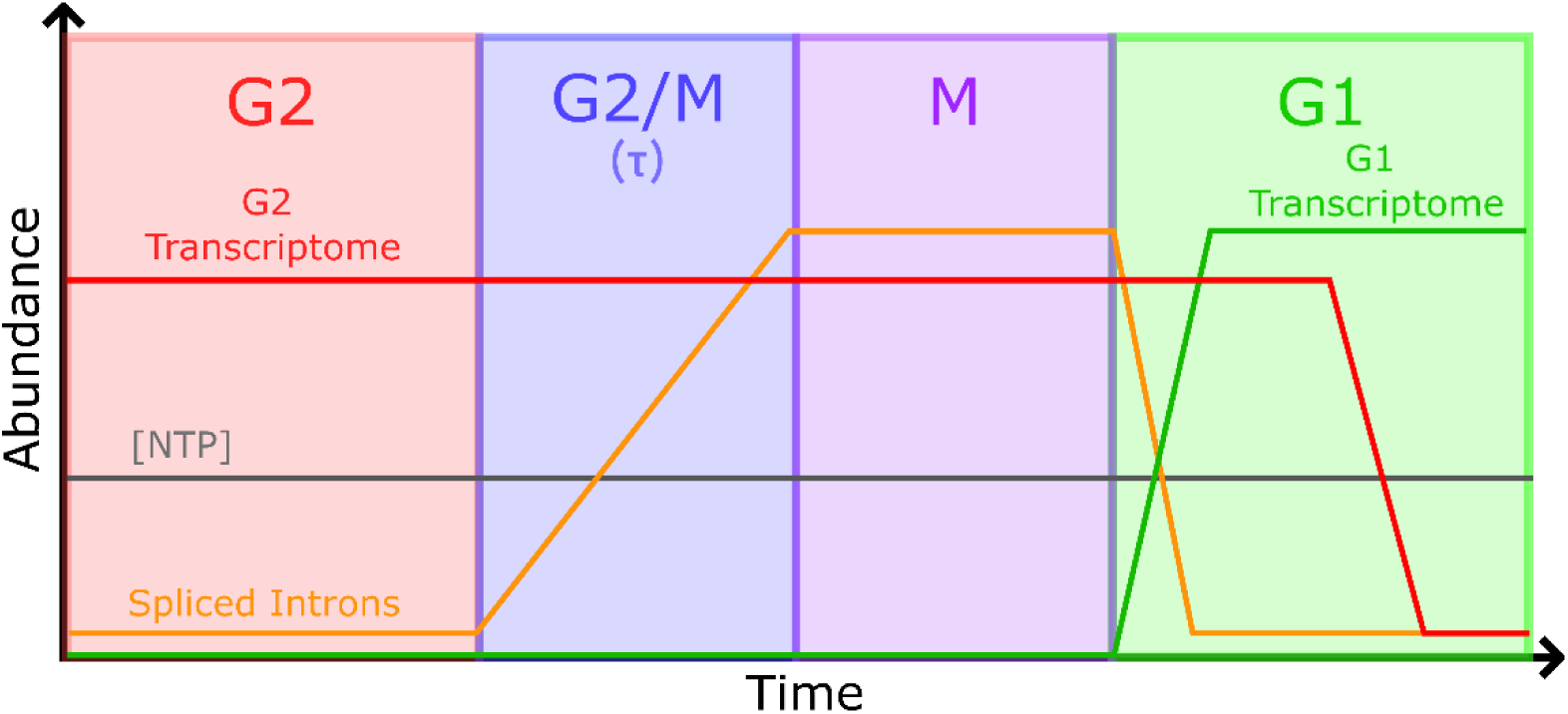
Diagram illustrating the hypothetical development of NTP concentration through mitosis (gray), G2 transcriptome (red), G1 transcriptome (green), and freely spliced introns (orange).

Evolutionary pressure is ignorant of immediate temporal dynamics. We must shift our mathematical frame of reference from an analytical kinetics perspective to an evolutionary optimization perspective. The pressure imposed by both the mitotic state and the G1 state will be incorporated into one determination of intron length needed over vast quantities of evolutionary time. From an iterative evolutionary modeling perspective, if a singular gene increases its expression profile in G1, then the cell would need to preserve more NTPs during τ, thus increasing *i*. Similarly, if gene expression is lowered during τ then pressure is exerted during G1 as a lack of NTPs and more intronic length is required. The question then is does the increased demand in intronic length necessarily need to be placed within the gene experiencing modified kinetics? If the system is otherwise static and unchanging, then no, it is not necessary. Many other genes could cover the burden of one iterative change. In this scenario the change could be reasonably integrated across all genes and minimal impact on the changing gene’s intronic content would be noticed. But gene expression is a hyper dimensional dynamic system. Small permutations will negatively permeate pressure though the transcriptome hyperspace. The direct comparison of summations is therefore not the valid way to determine our desired variable *i*. What we can determine is that by removing the summations, the pressure constraint imposed upon introns becomes 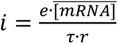 (Eq. 1).

In this hypothesis, Equation 1 is only a pressure, not a strict deterministic evaluation of intron length. Over vast quantities of evolutionary time, the pressures from the mitotic state and G1 sate will approximate Equation 1 in global dynamics, however, the system can still result in stochastically distributed lengths between genes to satisfy the overarching integration. Given there are over 20,000 genes (A.K.A. degrees of freedom) in the human genome for which the sum can be distributed, great variation in the individual gene-wise variance is to be expected. Furthermore, for complex multicellular organisms, Equation 1 would be optimized not only on one combination of G2/M and G1 transcriptomic states but all possible G2/M and G1 states across all possible cell types. Transcription is also stochastic (termed transcriptional bursting | (20)), making the contribution of a given gene dependent on the probability of binary state dynamics (on/off) of the gene bursting during the time τ (reference the distribution of *EERFI1* in Figure 2). For all these reasons, and likely a myriad more, the variance in *i* is expected to be unpredictable given the scale of the system. The trend aggregate mean around any evaluation of *i*, however, should match Equation 1 as the mean surrounding an evaluation of Equation 1 represents the reflection of global pressure applied weakly across those individual genes.

#### Derived Model Fits Transcriptional Kinetics Data in Mouse

The ability to accurately access the kinetic intron hypothesis’ quantitative predictions is greatly limited. There is only one reliable data set which reports the most critical pieces of data: mRNA synthesis rates, mRNA steady state abundance, and mRNA decay rate. Most literature data reports mRNA decay rate, some report mRNA synthesis rates, but coupling those measurements together with mRNA copy number to access the mRNA steady state relationship is strictly unique to Schwanhäusser et al. (19). To complicate matters, high throughput mRNA decay rates are not in wide agreement. Median/mean mRNA half-life data for human has been reported as median 10hr (21), mean 6.9hr (22), median 5.3hr (23, 24), 3.4hr (25), median ~0.8hr (26) in human. For mouse the half-life data has been reported as median 7.9hr (19), 3.6hr–5.4hr (24), 4.9hr (27), 2.9hr (28), or median of all RNA species 1.97hr (29). Many variables could be driving this irreproducibility between studies; notably, however, the data is reproducible within their individual studies. What this indicates is that using the steady state relationship with decay rates alone is inadequate, noting that Equation 1 could further be idealistically represented under certain assumptions as 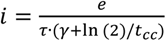 where γ is the decay rate and *t_cc_* is the cell cycle time (30).

To most accurately assess the model, data acquired from Schwanhäusser et al. enables the most robust analysis (19). The principle aim of their study was filling in the entire central dogma steady state models for both mRNa and protein dynamics: capturing the complete canonical DNA→mRNA→Protein paradigm. To do this they acquired mRNA steady state values (copy number per cell) and mRNA half-life measurements. Combining the two with a comprehensive cell cycle dependent steady state model they were able to determine mRNA synthesis rates. Importantly, normalizing by cell cycle state further normalizes synthesis rates demanded of Equation 1, noting that synthesis rates would be *r_G_*_2_ = 2 · *r_G_*_1_ in steady state assumptions due to the increase in loci in G2. As the most robust and complete publicly available data set it is the best available data to test the model. However, the mRNA half-lives used to calculate mRNA synthesis rates was on the higher end of reported half-lives range; with total mRNA half-life at 7.9hr for all RNAs or 9hr for those paired with their mRNA copy number and translation counterparts used in the analysis.

Matching the Schwanhäusser et al.’s data to the mouse genome mm10 (31), Equation 1 could be mostly satisfied. The exact kinetics between the G2/M checkpoint and the transcriptional silencing of DNA during prophase (τ) can be estimated by referencing CDK1 activity. Note, the duration must be at least long enough to produce the M transcriptomic state depicted in Figure 1. Considering CDK1 phosphorylation activity from FRET measurements, the time can be estimated between nuclear envelope break down at ~30min and metaphase at ~60min (32). This is approximately in line with the time it takes for CDK1-inhibtor RO-3306 synchronized cells to reach mitosis following release using a 45min mitotic shake off collection (33). Prior to this inhibition leading into mitosis, unspliced introns would have already accumulated at active transcription sites for a period of ~10min (11). In total I will estimate an intron lariat preservation time of 55min (45min RO-3306 shake off + 10 min prior accumulation) to describe the dynamics captured in Figure 2.

Analysis of Schwanhäusser et al.’s data set revealed a slope of ~1 and intercept ~0. Plots of the gene-wise calculations of Equation 1 in Fig. S2A shows a marginally poor Pearson coefficient of r=0.33 and spearman coefficient ρ=0.46 in the density scatter plot. Binning the data reveals the underlying global trend and hypothesized pressure constraint (Fig. S2B). It should be highly emphasized that while the linear fit and Pearson coefficient of the binned data reflect the linearity of the global trend, binning already correlated data will further emphasize the underlying correlation, decreasing the correlations relevancy to the evaluation of the model. Though the binned data is reflective of a characterization of the underlying pressure constraint imposed upon the system. Binning the data reveals the underlying global trend is exceptionally linear and that there exists an outliner near the far extreme of the binned data. Therefore, the extreme data was excluded from Fig S2B and values greater than 60,000bp were excluded in Fig. S2A accordingly, which accounted for 3.4% of the total data.

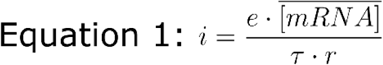

**Supplemental Figure 2:**
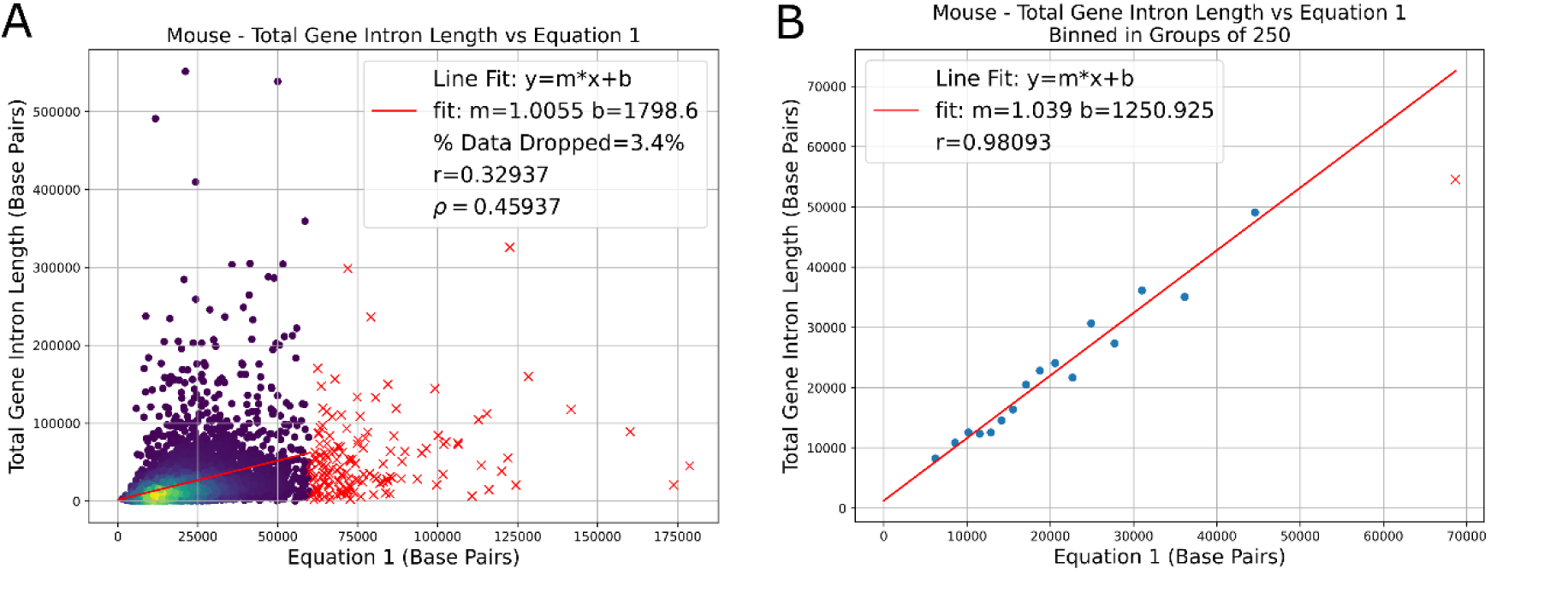
Evaluation of Equation 1 in mouse fibroblast cells using gene expression data from Schwanhäusser et al. and intron annotations from the mm10 mouse genome. (A) Gene-wise comparison of Equation 1 predictions as a function of total intronic length per gene. (B) Binned representation of the data in (A), where each point represents the mean of 250 genes, illustrating the global trend. Genes corresponding to outlier bins in (B) were excluded from the gene-wise representation shown in (A). ** Gene expression data were obtained from Schwanhäusser et al. (19), and intron length annotations were derived from the mm10 mouse genome annotation (17). All analyses and figure generation were performed in this study.

The existence of the extreme outliner was not surprising given the results in Figure 1 and Supplementary Table 1 which show extreme intron lengths would be most likely to occur during the mitotic transcriptomic state, for which this data set does not represent. The outliers can be expected following the inverse relationship in Equation 1 which shows 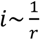 and since large introns represent the scenario where *r_interphase_* < *r_mitosis_* (gene expression increases during mitosis for intron dense gene | Figure 1) the predicted *i* using interphase values would be overestimate (*r_interphase_* > *r_mitosis_*). Table 1 further showed most genes were not greatly up or down regulated and as such the majority of genes should, and did, follow the model within the lower bound of predicted values. Supporting this, a greater spearman coefficient (ρ) as compared to Pearson coefficient reported between the two trends signifies that outliers may be impacting the evaluation of the Pearson coefficient. Furthermore, longer half-lives are less reliable than shorter half-lives as a limitation of pulse-chase labeling when fitting/extrapolating half-life measurements. Longer half-lives are also less accurate due to cell cycle dynamics disrupting steady state measurements. These longer half-lives are related to intron length via 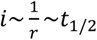, thereby also making larger intron calculations less reliable. To test if this influenced the fit, the data was iteratively tested against progressively limited upper thresholded half-life values [no threshold, 24hrs, 22hr, 20hr, 18hr, 16hr, 14hr, 12hr, 10hr, 8hr, and 6hr]. At and below the 10hr threshold the slope increased marginally to ~1.3 using ~50% of the data, but between no threshold and 12hr thresholds the slope remained unchanged at ~1. Such did not indicate concerns of experimentally confounded long half-life data biasing the measurements (as also assessed and discussed by Schwanhäusser et al. for their analyses).

It should be emphasized that RNA synthesis rates are the primary determinant of mRNA steady state; mRNA decay has been shown to not correlate greatly with steady state concentrations (19). Because synthesis rates were determined from steady state mRNA concentrations (#mRNA per cell), the correlations between intron length, mRNA steady state (Figure S4A and S4B), and synthesis rates (Figure S4C and S4D) cancel in Equation 1. Effectively, the correlation in Equation 1 is carried by the relationship between intron length and exon length (Figure S4E and S4F) multiplied by a factor of ~12. Showing that the Castillo-Davis et al. relationship (6) between mRNA abundance vs intron length can collapse in relationship to intron length vs exon length to produce a slope of 1 and intercept of ~0 from natural variables is still surprising, however.

### Highlights of the Existing Literature

#### Temporal Lag Between G1 Transcriptome Synthesis and M Transcriptome Degradation

Three key studies have discussed the kinetics of RNA degradation and RNA synthesis leading into G1. When comparing the studies it is critical to highlight differences in experimental methodology. The group studying RNA degradation synchronized cells with RO-3306 (33) while the groups studying RNA synthesis leading into G1 synchronized with nocodazole (14, 15). RO-3306 better maintains the kinetics of mitosis, as it is a CDK1 inhibitor which synchronizes cells to the G2/M checkpoint. Nocodazole is a microtubule destabilizer which prevents alpha tubulin from forming aligning condensed chromosomes into the metaphase plate. Release from nocodazole requires time for the alpha tubulin to polymerize and then subsequently form metaphase plate. Palozola et al. indicates vaguely that their metaphase occurs between 40 and 80 minutes post nocodazole release with early G1 occurring ~300 minutes post nocodazole wash-out (14). Hsiung et al. on the other hand indicate anaphase occurring 40 minutes post nocodazole washout with early G1 occurring between 40 minutes and 60 minutes post nocodazole (15). Since G1 in physiologically healthy mitosis should start 20 minutes after metaphase, these dynamics are clearly confounded by the nocodazole treatment. This critically changes the interpretation of the results when compared against one another.

The RNA degradation study by Krenning et al. revealed two waves of RNA degradation (33). Using pulse chase labeling, the first wave of mRNA decay comprised ~24% of RNAs by mitotic exit and the start of G1, 20 min post metaphase. The second wave of labeled mRNA was degraded much later into G1, 80 minutes post metaphase, and contained the bulk of the remaining transcripts. This postponed M phase transcriptome wide turnover and its kinetics aligns with the expectations of proposed model.

The synthesis kinetics of the G1 transcriptome post mitosis is complex and confounded between the two forementioned studies, but both studies agree that there is a ‘hyperactive’ burst of transcription at mitotic exist. The cleanest data was collected by Hsiung et al. (15) where by RNA Polymerase 2 (Pol2) loading and transcription reactivation was captured via ChIP-Seq on Pol2. There were two populations of genes: one which spiked at 60 minutes (corresponding to mitotic exist) and then plateau (~50% of genes) and the other loaded Pol2 more gradually over time (~38% of genes). Rather than quantifying Pol2 loading with ChIP Seq, Palozola et al. used pulse chase labeling to measure newly transcribed RNA transcripts. As previously mentioned, the reported timings of the mitotic index post nocodazole (recorded times of metaphase, anaphase, telophase, and early G1) varied greatly from that of Palozola et al. Furthermore, they used 40-minute periods of pulsed cell-permeable 5-ethynyluridine labeling to define Pol2 activity. This amount of time is insufficient for capturing the rapid progression of gene reactivation within healthy mitosis. In short, the two studies roughly validate each other, but Palozola et al.’s study does not seem to have the fidelity required to reasonably compare against Tanenbaum et al.’s RNA degradation observations. Ideally recollecting Hsiung et al.’s ChIP-Seq data using RO-3306—rather than nocodazole—would provide better comparative results.

From these studies it appears plausible that the G2 transcriptome NTP pool is decoupled from the G1 transcriptome NTP pool. Such could facilitate the need for introns as a NTP reservoir as proposed by the proposed model.

#### Spliceosomal Introns Emerged During Eukaryogenesis

Perhaps the largest question about introns is the following: Why are spliceosomal introns unique to eukaryotes? This hypothesis offers a very direct answer to the ancient question. Mitosis emerged during eukaryogenesis, and introns coevolved along with it. Spliceosomal, lariat derived, introns give control to the cell to modulate their turnover. The steps of eukaryogenesis were perhaps the most radical kinetic transformations to occur to life since the last universal common ancestor. The thinking is almost backwards to progress into a transient state where one shuts down the central dogma to divide. Such an action doubtlessly propagates a wave of kinetic mayhem, like the proposed notation that early G1 synthesis would demand a greater amount of NTPs than the NTP steady state could provide. Of all the things we can say about life, we can be assured maintaining an intracellular homeostasis is on that list. It seems reasonable then that a coordinated effort might have taken place—and continue to take place—to ensure homeostasis is maintained, even if the cost is as exorbitant as introns.

#### Saccharomyces cerevisiae Acts as a Negative Control

Notoriously, *S. cerevisiae* is known as being uniquely intron poor (34). A long-standing question is why? Lesser known is that S. cerevisiae undergoes only minor chromosome condensation during mitosis (35). A couple of authors have speculated that this laxer condensation during mitosis uniquely enables transcription to ensue during mitosis for S. cerevisiae (36, 37), though it should be noted that direct evidence is lacking. A literature search for mRNA transcription dynamics in *S. cerevisiae* during mitosis was only able to find one such examples of transcription during mitosis (38). It could also be that—due to the unique dynamics of budding—the distinction between G2 and M makes examining transcription during “mitosis” difficult to discern and study. Defining the onset of mitosis is not as easy as compared to canonical mitosis as a lot of mitotic processes occur simultaneously with the G2 phase. However, rDNA during mitosis is known to still be highly compacted like during canonical cell division and transcriptional silencing of that chromatin has been quantitatively assessed (39). This repression of rDNA and generally repression of RNA polymerase 1 during mitosis and early G1 is thought to be a general characteristic of rDNA expression in eukaryotes at large (40). In the context of The Kinetic Intron Hypothesis, the early G1 repression of rRNA synthesis may be taken as a step to limit competition for NTPs used for the synthesis of the G1 transcriptome. In S. cerevisiae though, the consensus appears to be that while rDNA is silenced, most genes are generally still transcribed though mitosis. This observation negates the functions of the proposed model since the G2/M transcriptome can transition continuously into G1 rather than undergoing the discontinuous canonical pathway. From this, introns would thus lack a purpose and be truncated and excised as a resource burden. That being, S. cerevisiae is the negative control species for the hypothetical model.

Like *S. cerevisiae*, multiple versions of mitosis have evolved. It could be that the differences between distant organisms and the comparative lengths of their introns are explainable through this view. Furthermore, mitochondria are also known to possess introns and could play a role in the model (41), though this is not the case for mitochondria in the kingdom animalia which do not contain introns. In many ways the model organisms of humans and mice might be an ideal system to evaluate the model due to their relatively straight forward mitosis dynamics and mitochondria genomes.

#### Rapidly Regulated Genes are Intron Poor

In 2008 Jeffares et al. published a paper titled: Rapidly Regulated Genes are Intron Poor (42). Simply enough, they found that genes which are regulated in response to stimuli or cell cycle changes have a suppressed intronic density. The finding was more prominent for yeasts (fungi) and thale cress (plant) while the correlations in mouse were weaker. Beyond the results of mitotically transcribed genes in Table 1 and Figure 2, this result is exactly in line with the expectations of the proposed hypothesis.

The kinetic intron hypothesis only demands the pressure for intron content to be within genes expressed leading into mitosis. Genes exclusively expressed at any other stage of the cell cycle or upon signal stimulation would be excluded from the applied pressure of Equation 1. To my knowledge, the only known published data set with precise mitotic gene expression profiles was by Tanenbaum et al. (7). Most studies report G2/M together as one measurement, which would further confound Jeffares et al.’s conclusion that mitosis is not linked to intron poor genes; especially since G2 is positively linked to intron poor genes in Figure 1. Likewise, genes responding to stimuli would not have a need for introns except for the more specialized benefits that having introns carry; such as NMD, splicing, or intron delay.

The study of intron less genes has reported similar findings. Performing GO analysis on intron less genes revealed overwhelmingly that 52.67% of intron less genes are involved in signal transduction activity (signal that changes the activity or state of the cell | (43)). The second largest classification was nucleic acid binding genes at 21.17%. While some key nucleic acid binding proteins are needed during mitosis, most proteins are strictly evicted from condensed chromatin. A lack of intron density in this subset of genes could therefore be reasonable within the context of the proposed model.

### Proposed Experiments

#### Validating Spliced Intron Accumulation During Mitosis

It is critical to this model that intron lariats persist into mitosis. The randomly selected set of FISH probes used here is simply not enough to firmly validate that the effect is global across all introns, though it induces skepticism towards the current paradigm. Preferably the use of Multiplexed Error Robust FISH (MERFISH) would resolve the order of magnitude necessary to make such claims (44). MERFISH can image 1,000s of mRNA species simultaneously and thus would be able to capture the dynamics of Figure 2 except across many different genes. Furthermore, by tracking a select set of known cell cycle dependent transcripts, the model can be tested with the utmost accuracy by creating a cell cycle pseudo time base on patterns of gene expression (45). The equality *N* = Σ *i_g_* · *r_g_* · τ would be directly observed as the number of intronic FISH species identified; the persistence of intronic RNA would necessarily come from the lead up until mitosis for the duration of τ (noting that τ is still not precisely known). The G1 state would furthermore be easily accessible to complete the full equality. Such would also replicate the preferential bias of intron length within mitotic genes presented in Figure 2.

A simpler experiment is to use the polyclonal intron-MS2 gene trap cell line created by Wan et al. (11). This cell line utilized a gene trap to insert MS2 stem loop arrays into intronic regions. Using these cells, the live cell dynamics of intron splicing were then studied across a population of genes simultaneously. Within the polyclonal cell line are 772 unique introns across 442 genes, with ideally one MS2 insert per cell. Since the goal is to visualize the persistence of many intronic species into mitosis, performing FISH on the MS2 sequences on the polyclonal cell line should reveal a heterogenous distribution of MS2 RNA species during mitosis. Not all genes are guaranteed to be transcribing—or rather transcriptionally bursting—during the lead up to mitosis, but a reasonable non-zero measurement across a random selection of cells would be a good indicator that the population of MS2 labeled intron species are universally persisting into miosis.

A third method to validate the persistence of intronic RNA into mitosis is through circular RNA sequencing. Hypothetically, the spliced introns should be in lariat form if the point of regulation occurs with DBR1 activity (discussed later on). It should be noted that circular RNA sequencing is different in nature to normal RNA sequencing. Due to their looped nature, lariat introns take on peculiar nonuniform sequencing patterns during a conventional RNA sequencing protocol. To overcome this, RPAD (RNase R Treatment, Polyadenylation, and Poly(A)+ RNA Depletion) was create to produce highly pure circular RNA extracts (46). The resulting solution should be all circular RNA. A cDNA library using random hexamers can then be acquired. It should be noted that preferably the RNA extract would come from a RO-3306 synchronized cell line post shake off, similar to how the G2/M/G1 RNASeq data was acquired by Tanenbaum et al. (7). Similar to this scheme, RNase R treatment on FISH samples can be used to validate intron circularity *in situ* using more refined methods (47).

#### Investigating the Effects of DBR1 During Mitosis

Debranchase (DBR1) is the leading culprit for intron lariat accumulation. As the rate limiting step in intron lariat turnover, a simple disruption to DBR1 activity could induce the effects seen in Figure 2. It has recently been shown that TTDN1 interacts with DBR1 to promote a ~19-fold increase in DBR1 catalytic activity (48). TTDN1—also known as M-Phase Specific PLK1 Interacting Protein—is known to be specifically phosphorylated and active during mitosis. During this time TTDN1·DBR1 interactions at DBR1’s c-terminus may be inhibited, thereby resulting in decreased catalytic turnover of intron lariats. To test this, a Förster resonance energy transfer experiment could be performed to determine the proximity of DBR1 to TTDN1 through the cell cycle. Another method would be through coimmunoprecipitation from mitosis synchronized cellular extract.

An interesting follow up experiment would be to perform crosslinking immunoprecipitation and sequencing (CLIP-seq) on DBR1. TTDN1 is thought to stabilize the lariate-DBR1 complex, thus promoting efficient debranching. It is questionable though, under this hypothesis, whether DBR1 itself possesses inhibitory mechanisms to bind the RNA. Pulling down DBR1 and sequencing any attached RNA species may point towards interesting mechanistic changes between the dynamics of RNA, DBR1, and TTDN1.

Another interesting feature to be explored involving DBR1 is cytoplasmic lariat debranching. As shown in Figure 2, free introns make their way into the cytoplasm after nuclear envelope break down. How these free introns get processed—assuming they are lariats—is of interest. There happens to be a suspicious phosphorylation site (S514) located directly adjacent to the known human DBR1 nuclear localization sequence (49–51). It has also been reported that this site is phosphorylated during G1 and mitosis. This suggests nuclear localization may be temporarily blocked, allowing DBR1 the ability to clear free intron lariats from the cytoplasm. To test this, immunofluorescence staining against DBR1 can be performed, looking for cytoplasmic occupancy of DBR1 during early G1. Comparing the kinetics of other nuclear protein shutting back to the nucleus post nuclear envelope reformation may reveal a temporal lag associated with DBR1 shuttling. A further method to test the S514 phosphorylation effect would be to use an expression vector—as used in the study to validate the nuclear localization sequence (51)—but with an aspartic acid or glutamic acid mutation at S514 to simulate the phosphorylated state.

### Observing NTP Abundance and Distribution Through Mitosis

To truly validate the workings of this model, an observation of NTP levels would be required at various points in time. It is specifically postulated that NTP levels are modulated by intron lariat turnover; or more precisely that NTP concentrations are constant and intron RNA promotes a momentary supplement during a NTP flux. The principal aim is to examine the observable of NTP concentration, or otherwise show that the population of NTPs within introns end up in the G1 transcriptome. Unfortunately, the observable of NTP concentration *in vivo* has been extraordinarily difficult to obtain reliably.

The ideal experiment would be to test if NTPs from introns end up within the G1 transcriptome. This is plausibly achievable via pulse chase labeling. Pulsing during the lead up to mitosis after RO-3306 synchronization would label newly synthesized intron RNA. In the follow up chase, the degraded labeled introns should act as a second pulse and transfer their labeled nucleotides to the newly synthesized G1 transcripts; thus, resulting in a double pulse. Notably, a homeostatic level of NTPs would exist which the labeled NTPs would mix with. A 1:1 incorporation would not be expected, but some ratio should transition into the newly synthesized pool. To complicate the experiment, s^4^U labeling during the pulse phase is very low. This experiment would need to maximally optimize the labeling efficiency, though this may be easier than normal pulse chase experiments as s^4^U toxicity to the cell may not be a concern; in total the experiment would last only ~2-4 hours with the pulse being only ~1 hour long. To avoid s^4^U toxicity, radio labeling could also be implemented at the cost of safety. One could also combined radio labeling pulse chase and s^4^U pulse chase, pulsing with radio labeling in G2/M then again with s^4^U to catch newly synthesized transcripts theoretically containing radio labeled NTPs. Such might also negate the effects of s^4^U toxicity while improving specificity. As a control, a theoretical DBR1 inhibitor could be used to block intron RNA turnover, thereby preventing the second pulse originating from intron lariats. New reports of a small molecule DBR1 inhibitor being discovered by Thomas Menees from University Missouri-Kansas City are circulating; though the inhibitor is weak and unlikely to facilitate the needs of the experiment (personal communications).

### Counter Argument

Many of the predictions of the kinetic intron hypothesis have yet to be examined. One avenue to critique, however, is the absolute abundance of NTPs inside cells. The spread of reported intracellular ribonucleic acid NTP levels varies greatly between reports (52), but the mean values have been reviewed and consolidated at being approximately 3,000uM for ATP, and 500uM for GTP, UTP, and CTP. To calculate the absolute number of NTPs, using BioNumbers reported HeLa cell volume of 2,425 µm^3^ (53), would put approximate absolute NTP values at 4.5 billion ATP, and 500 million GTP, UTP and CTP molecules. When calculating how many NTPs can be reserved in introns, taken from Schwanhäusser et al.’s data, would reveal approximately 200 million total NTPs or 50 million per NTP evenly distributed. ATP excluded, this rough calculation reveals that the reserved NTP concentration is an order of magnitude below the basal NTP levels. It is questionable therefor whether that lacking order of magnitude meaningfully could contribute to the greater whole.

This argument is cogent, but its oversimplifications also produces skepticism. It could be that the absolute number of NTPs is further distributed and in flux. Nucleotides exist in multiple states, such as mono-, di-, and tri-nucleotides. Kinetic transfer between the three for specific use is further facilitated by enzymes. Each further contributes functionality within their own identity, with only ribonucleotide triphosphates participating RNA synthesis. It is difficult to say whether the absolute order of magnitude difference is the actual difference when considering the system as a whole using nucleotides at optimized concentrations, the various mono-, di-, and tri-nucleotide distributions, and the presence of a second order flux of the system during the non-steady state dynamics of mitosis. The argument stands, however, that the magnitude of The Kinetic Intron Hypothesis may not constitute a large enough effect to buffer levels beyond steady state basal levels.

## Supplemental Tables and Figures

**Table S1.**
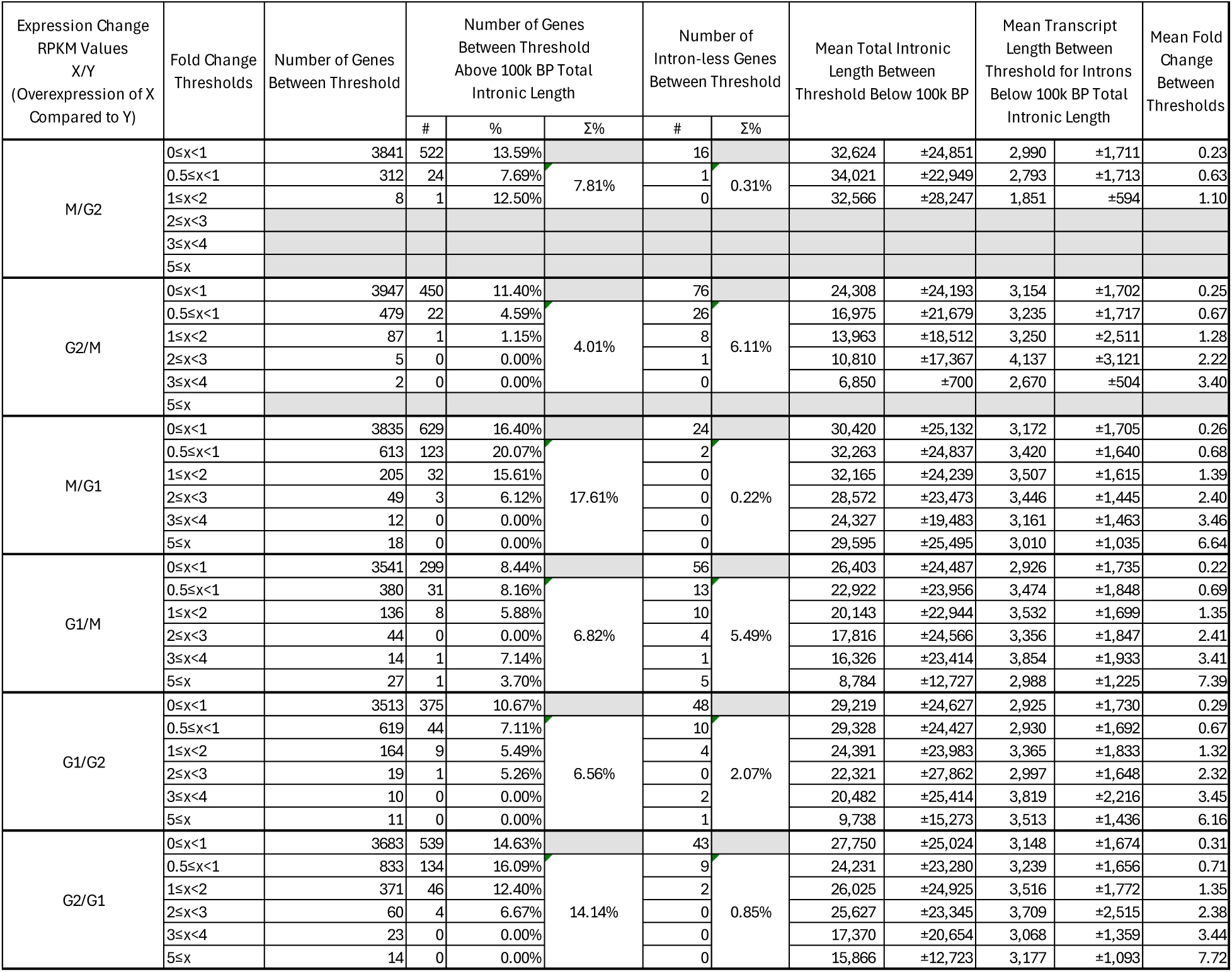
Comparison of intron length distributions across M, G1, and G2 transcriptomic states. **Gene expression data were obtained from Tanenbaum et al. (7), and intron annotations were derived from the hg19 human genome annotation (8).

**Supplemental Figure 3:**
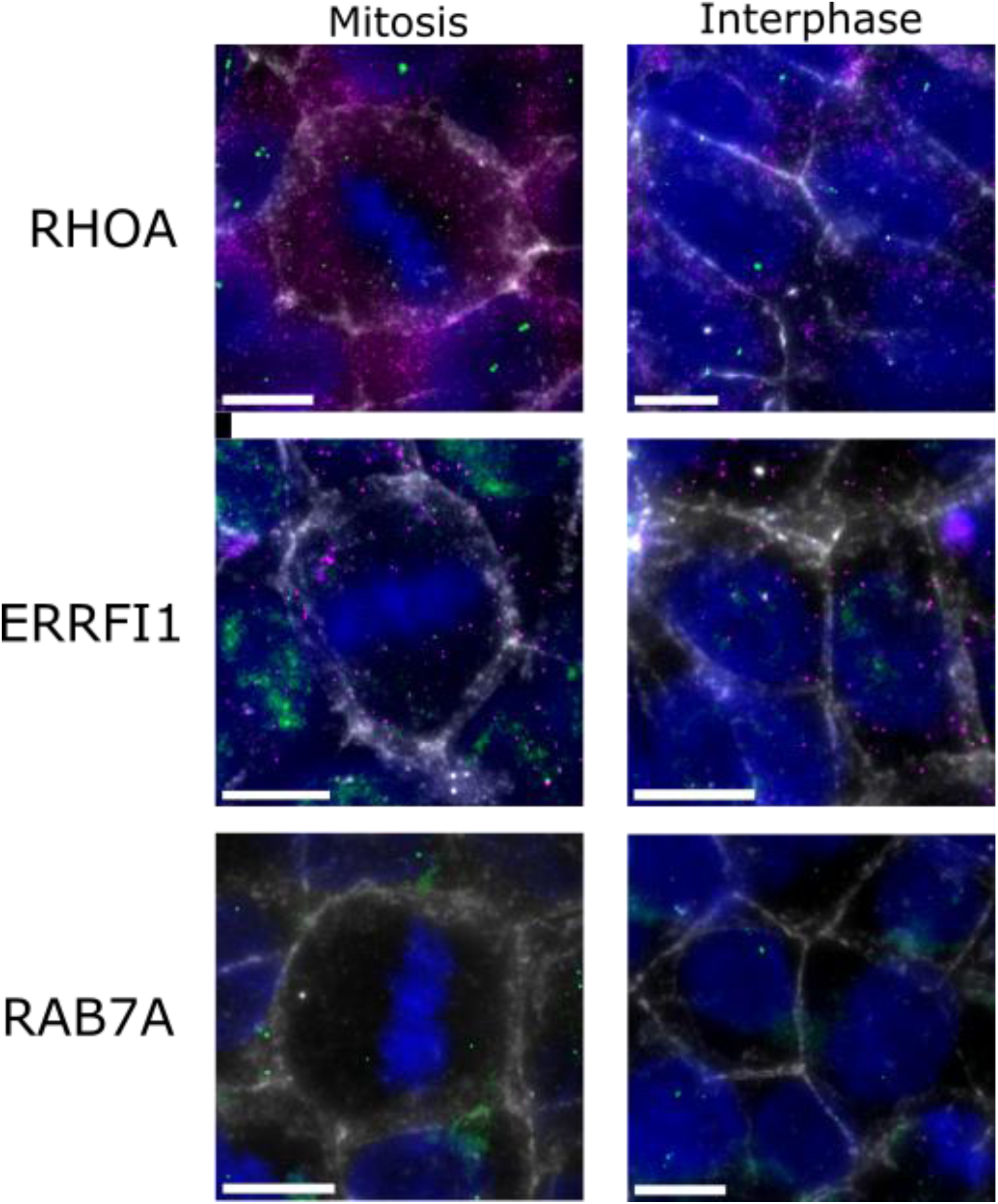
Examples of Figure 2 replicate using FISH-IF. Introns are shown in green, mRNA in magenta, E-Cadherin membrane IF in white, and DAPI in blue. Scale bars = 10um.

**Supplemental Figure 4:**
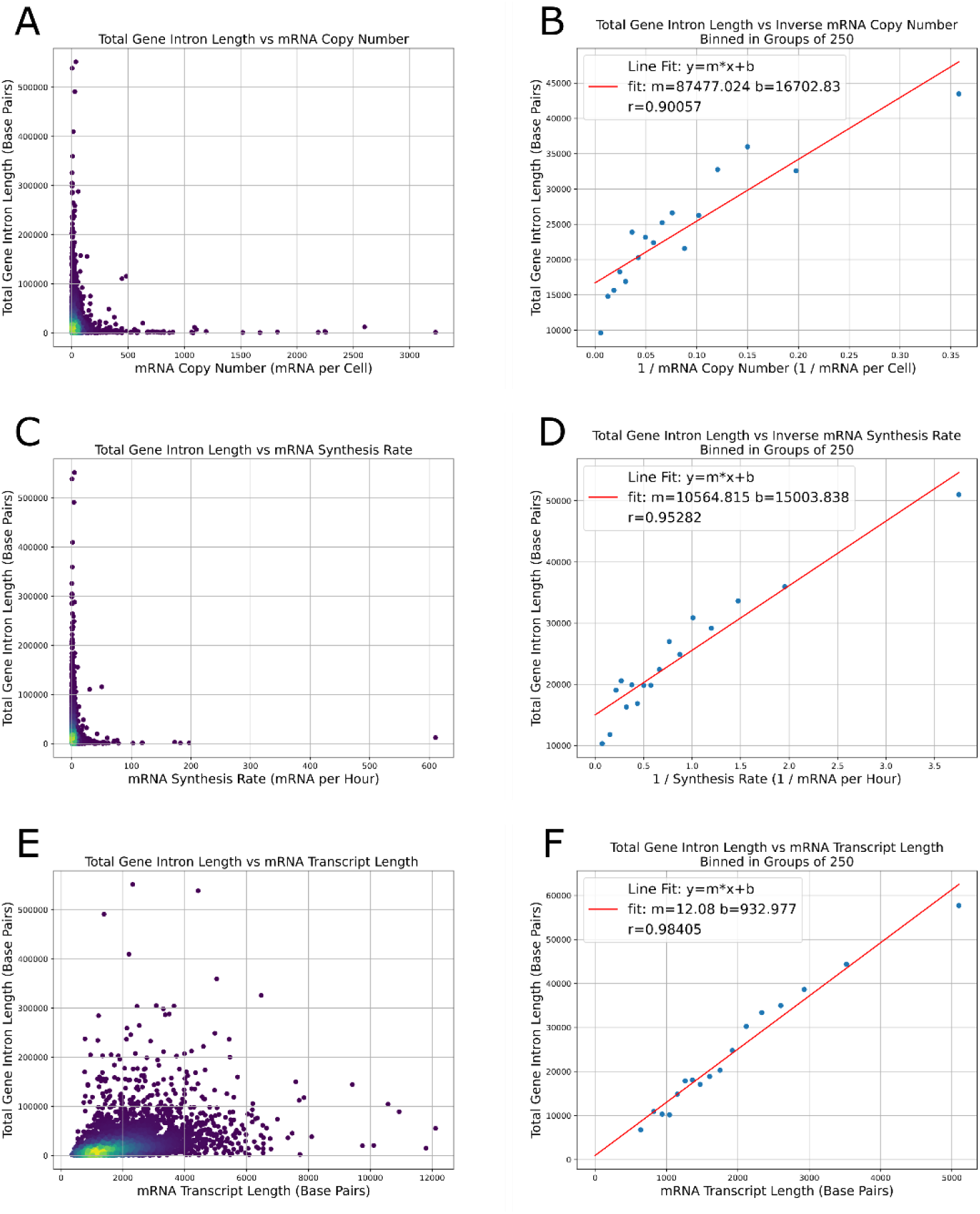
Scatter plots showing the trends of the variables used in Equation 1 against total gene intron length. Total intron length vs copy number is shown in A and inversed and binned in B to show the trend is inverse-like. Similarly, total intron length vs RNA synthesis rate is plotted in C and D accordingly. Total gene intron length and mRNA transcript length and its binned values to indicate global trends are further plotted in E and F accordingly. A and C reproduce trends previously reported by Castillo-Davis et al. (6). **Gene expression and RNA synthesis rate data were obtained from Schwanhäusser et al. (19), and intron and transcript length annotations were derived from the mm10 mouse genome annotation (17). All analyses were performed in this study.

## Notes

https://drive.google.com/drive/folders/1rT_KonGHGtawlSYYJnEgUVWjt-yqOUvZ?usp=sharing

### Competing Interest Statement

The authors have declared no competing interest.

